# The P681H mutation in the Spike glycoprotein confers Type I interferon resistance in the SARS-CoV-2 alpha (B.1.1.7) variant

**DOI:** 10.1101/2021.11.09.467693

**Authors:** Maria Jose Lista, Helena Winstone, Harry D Wilson, Adam Dyer, Suzanne Pickering, Rui Pedro Galao, Giuditta De Lorenzo, Vanessa M. Cowton, Wilhelm Furnon, Nicolas Suarez, Richard Orton, Massimo Palmarini, Arvind H. Patel, Luke Snell, Gaia Nebbia, Chad Swanson, Stuart J D Neil

## Abstract

Variants of concern (VOCs) of severe acute respiratory syndrome coronavirus type-2 (SARS-CoV-2) threaten the global response to the COVID-19 pandemic. The alpha (B.1.1.7) variant appeared in the UK became dominant in Europe and North America in early 2021. The Spike glycoprotein of alpha has acquired a number mutations including the P681H mutation in the polybasic cleavage site that has been suggested to enhance Spike cleavage. Here, we show that the alpha Spike protein confers a level of resistance to the effects of interferon-β (IFNβ) in lung epithelial cells. This correlates with resistance to restriction mediated by interferon-induced transmembrane protein-2 (IFITM2) and a pronounced infection enhancement by IFITM3. Furthermore, the P681H mutation is necessary for comparative resistance to IFNβ in a molecularly cloned SARS-CoV-2 encoding alpha Spike. Overall, we suggest that in addition to adaptive immune escape, mutations associated with VOCs also confer replication advantage through adaptation to resist innate immunity.

## INTRODUCTION

Both SARS-CoV-1 and SARS-CoV-2 enter target cells through the interaction of their Spike proteins with the angiotensin converting enzyme 2 (ACE2) cell surface receptor. Upon attachment and uptake, the Spike glycoprotein trimer is cleaved by cellular proteases such as cathepsins and TMPRSS family members at two positions – the S1/S2 junction and the S2’ site – to facilitate the activation of the fusion mechanism. Similar to more distantly related beta-CoVs, but so far unique in known Sarbecoviruses, the SARS-CoV-2 glycoprotein contains a polybasic furin cleavage site (FCS) with a (681-PRRAR-685) sequence at the S1/S2 junction. This allows the Spike precursor to be processed to the S1 and S2 subunits by furin-like proteases before viral release from the previously infected cell(Hoffmann et al., 2020). This leads to a proportion of processed Spikes to be present on the virion before engagement with the target cell, allowing for rapid activation and fusion at or near the cell surface by TMPRSS2. The importance of the FCS is highlighted by the observations that it enhances SARS-CoV-2 replication specifically in airway epithelial cells and is essential for efficient transmission in animal models (Peacock et al., 2021a).

The alpha variant of SARS-CoV-2 arose in the South-East of England in autumn 2020, and rapidly spread across the world in the first months of 2021. Various studies suggested that alpha had an increased transmissibility between individuals(Lindstrom et al., 2021; Mok et al., 2021; Tanaka et al., 2021). Alpha contains nine amino acid residue changes in Spike including a deletion of amino acid residues H and V in the N-terminal domain at position 69/70, thought to increase Spike incorporation into virions, a single amino acid deletion of Y144 (thought to assist NTD antibody neutralization escape), and a N501Y mutation in the RBD which enhances ACE2 binding affinity (Meng et al., 2021),(Chi et al., 2020). Together these changes have been shown to reduce efficiency of neutralization by some antibodies(Graham et al., 2021). Alpha also acquired a P681H change in the FCS which has been shown to increase the accessibility of the site by furin leading to enhanced cleavage(Mohammad et al., 2021),(Zhang et al., 2021). The alpha variant Spike has also recently been reported to mediate more efficient cell-to-cell fusion and syncytia formation (Michael Rajah et al., 2021),(Sanches et al., 2021).

We and others have previously found that SARS-CoV-2 is variably sensitive to entry inhibition by the interferon-regulated IFITM family (Winstone et al., 2021),(Shi et al., 2021). the three members of the family form multimeric complexes and have antiviral activity against diverse enveloped viruses by blocking fusion of the viral and cellular membranes (Bailey et al., 2014; Shi et al., 2017). While IFITM1 localizes primarily to the plasma membrane, IFITM2 and IFITM3 are internalized via a conserved YxxΦ endocytic motif to occupy distinct and overlapping endosomal compartments(Jia et al., 2012; Jia et al., 2014). The sensitivity of a given virus to individual IFITM proteins is largely determined by their route of cellular entry. We have previously shown that for a prototypic Wuhan-like SARS-CoV-2 isolate from early 2020, IFITM2 reduced viral entry and contributed to type I interferon (IFN-I)-induced inhibition in human cells(Winstone et al., 2021). Sensitivity to IFITM2 could be markedly enhanced by deletion of the FCS, suggesting that furin processing ameliorated SARS-CoV-2 sensitivity to IFITM2-restriction at least to some extent. We therefore postulated that the altered cleavage site of alpha may have consequences for its sensitivity to IFN-I and IFITMs. Here, we demonstrate that the Spike of the alpha variant is less sensitive to restriction by IFNβ and IFITMs in A549-ACE2 and Calu-3 cells. Furthermore, this resistance correlates with the enhanced polybasic site as reversion of this cleavage site increases the alpha variant’s sensitivity to IFITM restriction. Finally, we demonstrate that the H681P reversion in the full-length virus confers IFNβ sensitivity to alpha and suggest that part of this phenotype is driven by IFITMs.

## RESULTS

### The Spike proteins of currently circulating variants display differing sensitivities to IFITMs in A549-ACE2 cells

Previously we have shown that viral entry mediated by the original Wuhan-1 Spike pseudotyped lentiviral particle (PLV) or the England 02 isolate (hCoV-19/England/02/2020) was inhibited by IFITM2 in A549-ACE2 cells, and that this effect correlated in part with the IFNβ sensitivity of the virus(Winstone et al., 2021). Over 2020 and 2021 several major variants of concern (VOCs) have arisen – alpha (B.1.1.7) in the UK, beta (B.1.351) in South Africa, gamma (P1) in Brazil, and delta (B.1.617.2) in India. We wanted to compare the sensitivity of viral entry of the alpha, beta, gamma and delta to the presence of IFITM proteins, given that these variants have several changes in the Spike sequence (Figure 1A). Initially, PLVs bearing the spike protein of each variant to test whether they were restricted by IFITMs overexpressed on A549-ACE2 cells (Figure 1B-G). First, we confirmed that the D614G mutation in the parental Wuhan-1 Spike that became prevalent in the first wave of the pandemic displays a similar IFITM phenotype to the previously characterised SARS-CoV-2 Spike (Wuhan-1) (Figure 1B, 1C)(Winstone et al., 2021). The addition of D614G to the Wuhan-1 Spike had no effect on IFITM1 and IFITM2 sensitivity of PLV entry, while we observe a slight enhancement in the presence of IFITM3. We next compared the IFITM sensitivities of the major global VOCs. The alpha Spike appeared completely insensitive to IFITMs 1, 2 or 3 whilst beta, gamma and delta still retain some sensitivity to IFITMs 1 and/or 2 (Figure 1B-G). None of the VOCs were restricted by IFITM3 and, interestingly, we noted that IFITM3 appeared to markedly enhance entry mediated by the alpha variant Spike. Next, we pre-treated A549-ACE2-IFITM3 cells with cyclosporin H as this compound is known to drive IFITM3 to degration(Petrillo et al., 2018), and showed that this led to a specific abolishment of the enhanced infection by alpha PLVs (Supplemental Figure 1). Henceforth, we selected alpha to further investigate and determine the mechanism of IFITM resistance.

**Figure 1.**
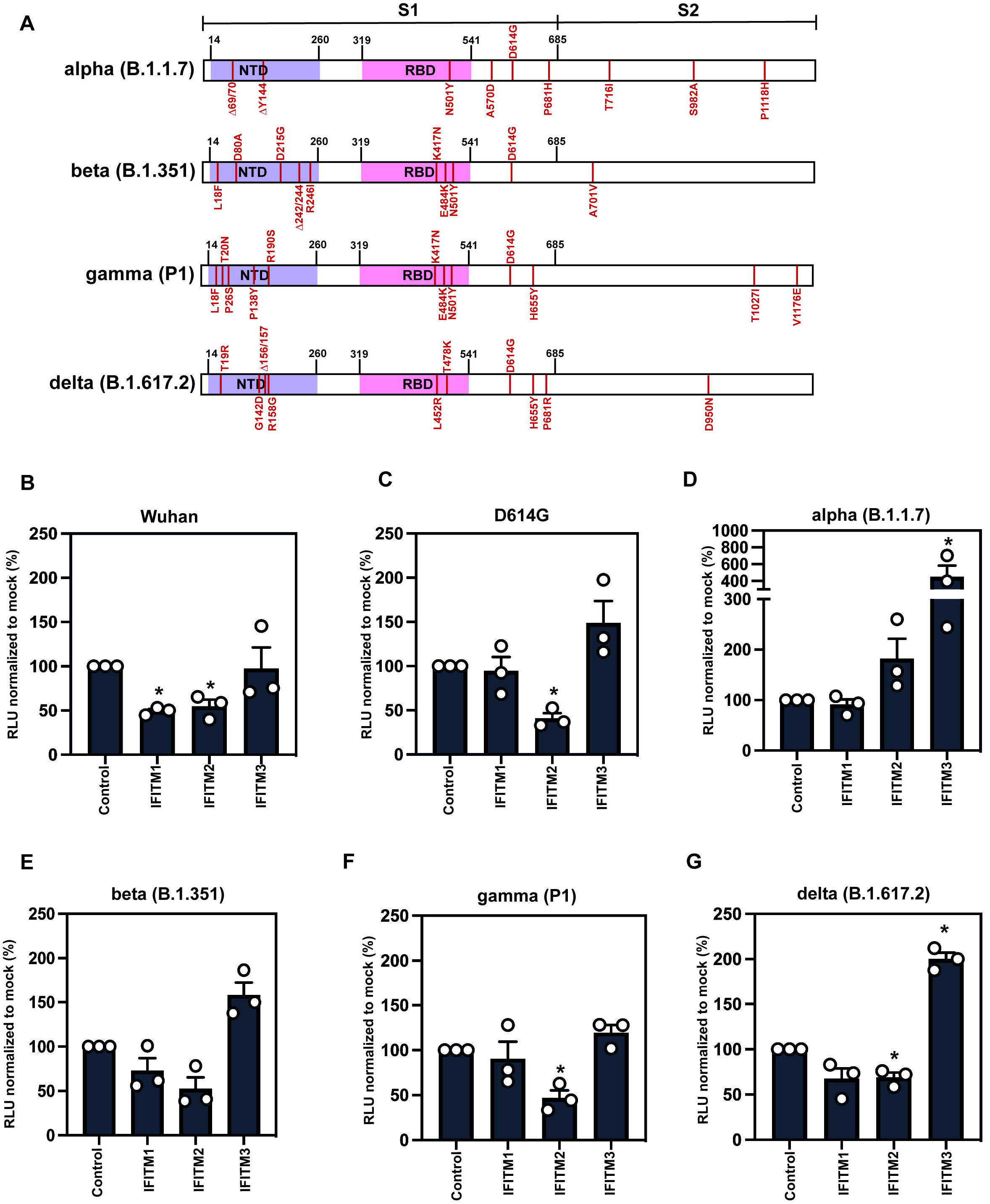
IFITM sensitivity of SARS-CoV-2 variants of concern. A) Schematic of Spike protein domains of the different variants of concern relative to the original Wuhan Spike sequence: alpha, beta, gamma and delta. The different mutations between the variants are represented in red. B-G) IFITM sensitivity of Wuhan, D614G, alpha, beta, gamma and delta PLVs in A549-ACE2 cells stably expressing the individual IFITMs. PLV entry was quantified by Luciferase activity 48 hours after infection and normalized to control cells. Data shown are mean ± SEM, n=3. Statistics were calculated in Prism using *t*-test, stars indiciate significance between control cell and individual IFITM (*P=0.05).

Next, to confirm whether IFITM sensitivity results seen with the alpha spike PLVs could be recapitulated with the full-length virus (Figure 2A), we infected A549-ACE2 cells (stably expressing IFITMs or not) with either the Wuhan-like England/02 isolate or alpha at an MOI of 0.01 and measured infection by qPCR of E copies in the infected cells 48 hours later (Figure 2B). As expected, replication of England/02 was significantly reduced in IFITM2 expressing cells over 48h. By contrast alpha replicated as well in IFITM2-expressing cells and to a significantly higher level in IFITM3-expressing cells in comparison to the control A549-ACE2 cells. This confirmed that alpha SARS-CoV-2 is resistant to the effects of IFITMs 1 and 2 and enhanced by IFITM3 on viral entry. Previous reports have suggested that the alpha spike is more efficiently cleaved than Wuhan-like isolates due to the presence of the P681H mutation optimising the accessibility of the FCS (Mohammad et al., 2021; Zhang et al., 2021) We immunoblotted our viral stocks of England-02 and alpha and confirmed that the alpha has more processed spike on the virions (Figure 2C, right panel), although it was not as clearly discernible a difference in processing on PLVs (Figure 2C, left panel). However, PLV infection in A549-ACE2 cells pre-treated with E64D, a cathepsin inhibitor, that inhibits the entry of SARS-CoV-2 in endosomal compartments because it cleaves S1/S2 junctions that have not been processed by furin. This showed that the alpha variant Spike mediated entry of PLVs is markedly less sensitive to this endosomal protease inhibition, when compared with the Wuhan-1/D614G Spike, indicating differences in the site of virus entry mediated consistent with enhanced Spike processing by furin in the producer cell (Supplementary Figure 2).

**Figure 2.**
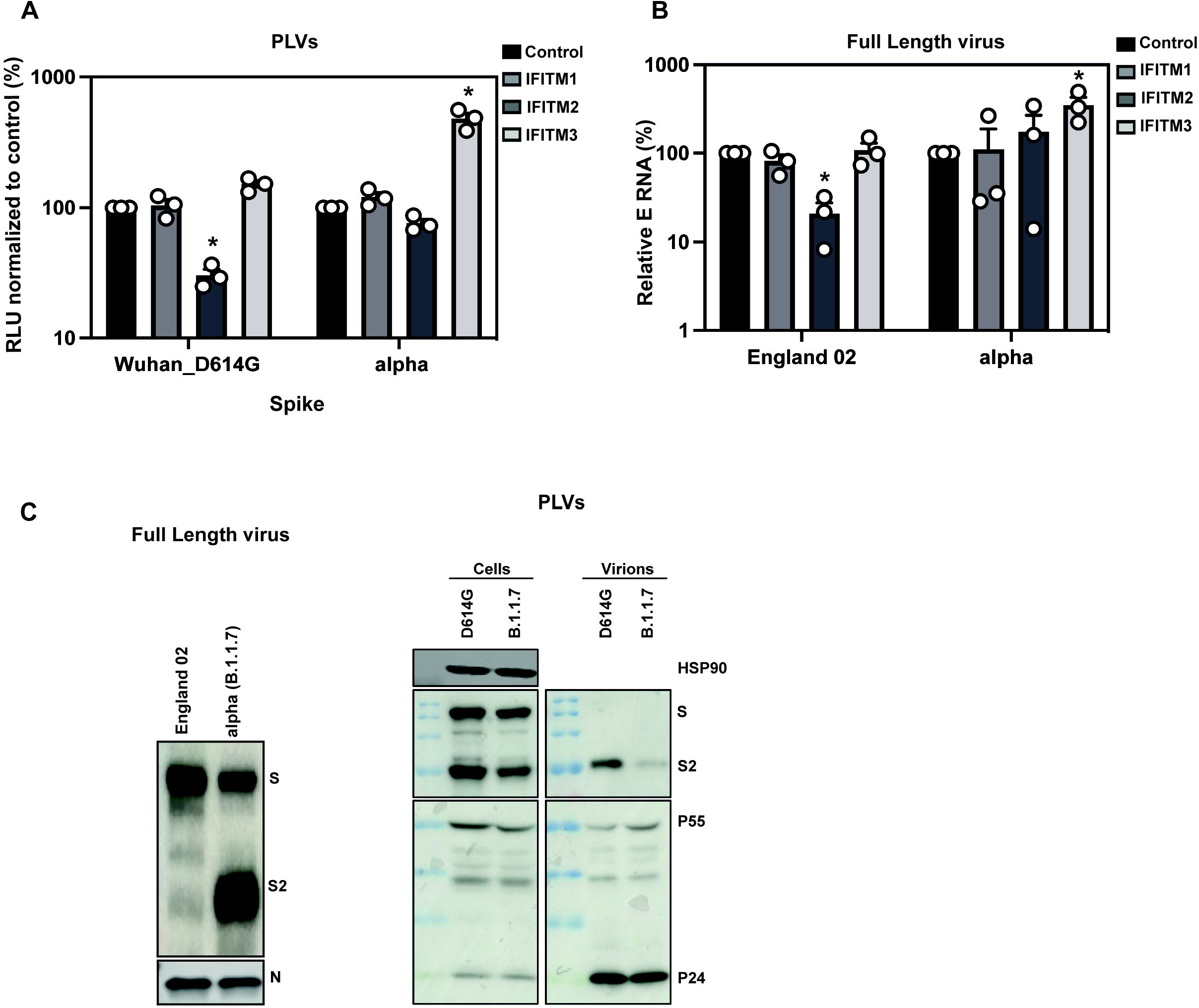
The alpha variant of SARS-CoV-2 is resistant to IFITMs. A) D614G and Alpha PLVs infection of A549-ACE2 cells stably expressing the individual IFITMs. Infection was quantified by Luciferase activity 48 hours later and normalized to control cells. Data shown are mean ± SEM, n=3. Statistics were calculated in Prism using *t*-test, stars indiciate significance between control cell and individual IFITM (*P=<0.05). B) Infection of A549-ACE2 stably expressing the individual IFITMs with England 02 and alpha full-length viruses at MOI 0.01. Infection was quantified by RT-qPCR of E gene relative to GAPDH 48 hours later; graph represents E mRNA levels relative to GAPDH. Data shown are mean ± SEM, n=3. Statistics were calculated in Prism using *t*-test, stars indiciate significance between control cell and individual IFITM (*P=<0.05). C) Western blot from representative D614G and alpha PLVs produced in HEK293T/17 cells, and virions from full-length England-02 and alpha viruses. Virions were purified through a 20% sucrose gradient.

### The alpha variant is less sensitive to IFNβ than a wave 1 isolate

While previous data has indicated that the original Wuhan-like SARS-CoV-2 virus can delay pattern recognition of viral RNA in target cells, its replication highly sensitive to exogenous interferon treatment in culture, in part determined by IFITM2(Jouvenet, 2021). Having confirmed that the alpha variant is resistant to IFITMs expression when ectopically expressed in cells, we then tested if alpha is also resistant to the effects of IFNβ, as suggested(Guo et al., 2021; Thorne et al., 2021). Indeed, we found from measuring supernatant viral RNA 48 hours after infection of both A549-ACE2 cells and the human lung epithelial cell line Calu3, which more faithfully represent target cells in the respiratory tract, that alpha is more resistant than England/02 to pre-treatment with increasing doses of IFNβ (Figure 3A, 3B). We have also confirmed this phenotype by measuring viral RNA in cell lysates, and further extending these obervations to two clinical isolates of alpha isolated (clinical isolates 10 and 28; Figure 3C). Thus in comparison to a representative example of Wuhan-1-like SARS-CoV-2, the alpha variant has a marked resistance to type I interferon.

**Figure 3.**
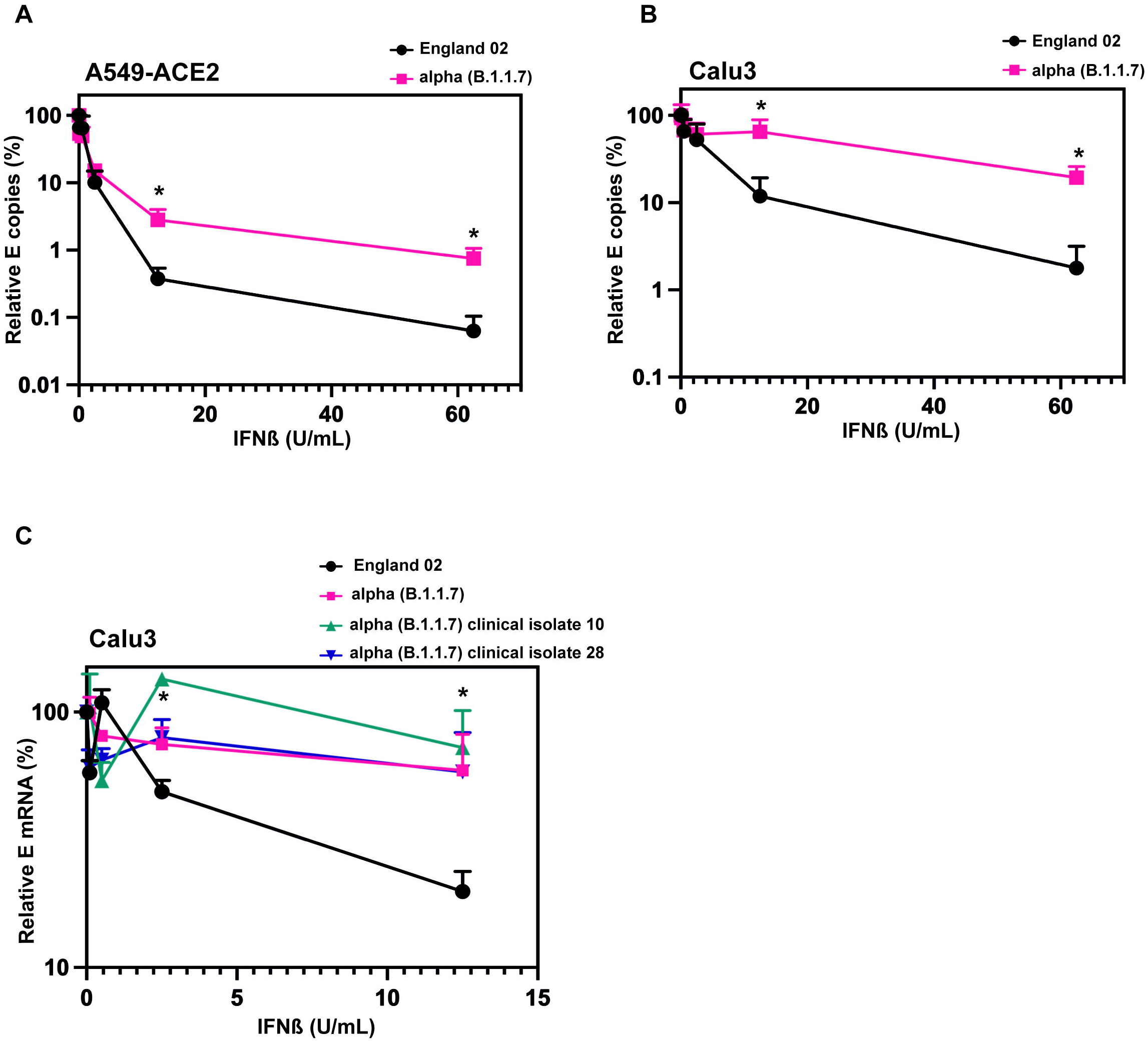
The alpha variant is resistant to IFNβ. A) England 02 and alpha full-length virus infection in A549-ACE2 cells pre-treated with IFNβ. Cells were pre-treated with increasing concentrations of IFNβ for 18 hours prior to infection with either virus at 500 E mRNA copies/cell. Infection was quantified by RT-qPCR of E mRNA from the supernatant 48 hours later and normalised to the un-treated control. Data shown are mean ± SEM, n=3. Statistics were calculated in Prism using *t*-test, stars indicate signifincance between the different viruses at individual IFN concentrations (*P=<0.05). B) England 02 and alpha full-length virus infection in Calu-3 cells pre-treated with IFNβ. Cells were pre-treated with increasing concentrations of IFNβ for 18 hours prior infection with either virus at 5000 E copies/cell. Infection was quantified by RT-qPCR of E mRNA from the supernatant 48 hours later and normalised to the un-treated control. Data shown are mean ± SEM, n=3. Statistics were calculated in Prism using *t*-test, stars indicate signifincance between the different viruses at individual IFN concentrations (*P=<0.05). C) England 02 and clinical isolates of alpha full-length virus infection in Calu-3 cells pre-treated with IFNβ and harvested as in A and B. Cells were pre-treated with increasing concentrations of IFNβ for 18 hours prior to infection with either virus at 5000 E copies/cell. Infection was quantified by RT-qPCR of cellular E mRNA relative to GAPDH 48 hours later and normalised to the un-treated control. Data shown are mean ± SEM, n=3. Statistics were calculated in Prism using *t*-test, stars indicate signifincance between the different viruses at individual IFN concentrations (*P=<0.05).

### The P681H mutation is necessary for conferring IFITM and IFNβ resistance in alpha

Our previous data indicated IFITM sensitivity of SARS-CoV-2 Spike can be increased by deleting the FCS. Given that alpha and delta Spikes have acquired mutations at P681 to H or R respectively, and these enhance Spike cleavage to S1/S2 (Supplementary figure 3), we hypothesized that P681H might be a determinant of resistance to IFN and IFITM. First we confirmed that none of the other individual alpha-Spike defining mutations were sufficient to confer IFITM resistance to a Wuhan/D614G Spike in PLVs (Supplementary figure 4 A–F). By contrast a P681H mutation in the Wuhan-1/D614G Spike was sufficient to abolish IFITM2-mediated inhibition PLVs entry, but not to fully confer the IFITM3-mediated enhancement phenotype (Figure 4A). As expected, deletion of the HRRA cleavage site in the alpha Spike conferred potent sensitivity to IFITM2 and reduced the level of enhancement we showed with IFITM3. Finally, we reverted the mutation in alpha to H681P and showed it gained IFITM2 sensitivity. Thus, the H681P mutation in the alpha spike confers IFITM resistance consistent with its enhanced furin cleavage. However delta bears a P681R mutation yet is not fully IFITM resistant (Figure 1G), implying that the P681H phenotype may be context dependent. Indeed, we found that combining the P681H mutation in the Wuhan spike with the deletion at position 69-70 in the NTD found in the alpha variant but not delta was sufficient to fully confer an alpha spike phenotype to D614G (Supplementary figure 4G). This suggests that the H681 confers IFITM resistance in the context of other adaptations in the alpha Spike that are thought to affect the conformation of S1 and the interaction with ACE2.

**Figure 4.**
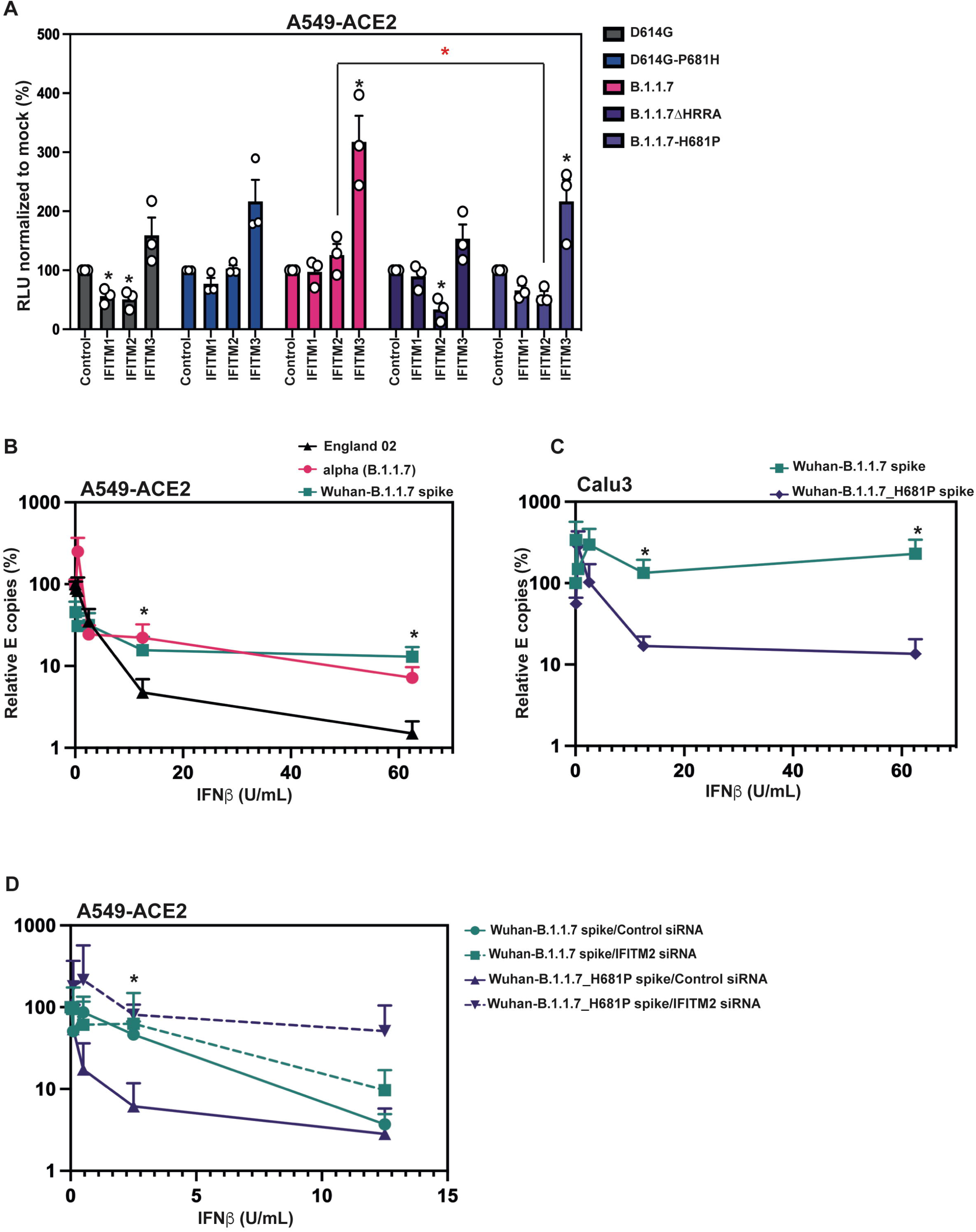
The P681H mutation is necessary but not sufficient for IFITM resistance, and necessary and sufficient for IFNβ resistance. A) D614G, D614G-P681H, alpha, alpha-ΔHRRA, and alpha-H681P PLVs infection in A549-ACE2 cells stably expressing the individual IFITMs. PLVs entry was quantified by Luciferase activity 48 hours later and normalized to control cells. Data shown are mean ± SEM, n=3. Statistics were calculated in Prism using t-test, black stars indicate significance relative to the control cells, red stars indicate significance between alpha and alpha-H681P in IFITM2 cells (*P=<0.05). B) England 02, alpha, and Wuhan-alpha Spike full-length virus infection in A549-ACE2 cells pre-treated with IFNβ. Cells were pre-treated with increasing concentrations of IFNβ for 18 hours prior to infection with either virus at 500 E copies/cell. Infection was quantified by RT-qPCR of E mRNA in the supernatant 48 hours later and normalised to the un-treated control. Data shown are mean ± SEM, n=3. Statistics were calculated in Prism using *t*-test, stars indicate signifincance between the different viruses at individual IFN concentrations (*P=<0.05). C) Wuhan(B.1.1.7 spike) and Wuhan(B1.1.7 spike H681P) Spike full-length virus infection in Calu-3 cells pre-treated with IFNβ. Cells were pre-treated with increasing concentrations of IFNβ for 18 hours prior to infection with either virus at 5000 E copies/cell. Infection was quantified by RT-qPCR of E mRNA in the supernatant 48 hours later and normalised to the un-treated control. Data shown are mean ± SEM, n=3. Statistics were calculated in Prism using *t*-test, stars indicate signifincance between the different viruses at individual IFN concentrations (*P=<0.05). D) A549-ACE2 cells were transfected with siRNAs against non-targeting control or IFITM2 for 24 hours and then treated with IFNβ for 18 hours prior to infection with Wuhan(B.1.1.7 spike) or Wuhan(B.1.1.7 spike H681P) at 500 copies/cell. Infection was quantified by RT-qPCR of E gene relative to GAPDH 48 hours later; graph represents E mRNA levels relative to GAPDH. Data shown are mean ± SEM, n=3. Statistics were calculated in Prism using *t*-test, stars indicate signifincance between the different viruses at individual IFN concentrations (*P=<0.05).

Having established that P681H change is necessary for the resistance of the alpha variant Spike to IFITMs, we next wanted to address if this was also a determinant for the resistance to type-I IFN resistance of the virus itself. We constructed a recombinant molecular clone of SARS-CoV-2 Wuhan-1 encoding Spike from the alpha variant. This virus essentially mimicked the resistance of the alpha variant itself to IFNβ in comparison to a representative Wuhan-1 like virus, England-02, demonstrating that the alpha Spike alone is sufficient to confer a level type I IFN resistance in A549-ACE2 cells (Figure 4B). We then took this recombinant virus and reverted the amino acid residue H681 to a proline. Importantly, this single point mutation was sufficient to confer a significant sensitivity to IFNβ in Calu3 cells (Figure 4C). Lastly, we wanted to confirm whether siRNA knockdown of IFITM2 was sufficient to rescue the IFN sensitivity of the Wuhan(B.1.1.7 Spike H681P) virus. We therefore knocked down IFITM2 expression using siRNA in A549-ACE2 cells and treated the cells with IFNβ as before. We found that the H681P reverted virus was rescued from IFNβ restriction during IFITM2 knockdown, meanwhile the Wuhan(B.1.1.7 Spike) virus was unaffected, consistent with this virus being resistant to IFITM2 restriction. Thus, this confirmed that the Spike protein of the alpha variant of SARS-CoV-2 is a determinant of type-I IFN resistance and that the P681H mutation is necessary for this.

## DISCUSSION

Here we have shown that the Spike protein of the alpha variant of SARS-CoV-2 is resistant to IFN-I. Furthermore, we show that this maps to the histidine residue adjacent to the FCS that has been mutated from the parental proline, which has been shown to enhance Spike cleavage at the S1/S2 boundary (Peacock et al., 2021b). This residue is necessary to confer resistance to IFITM2 and enhancement by IFITM3, and as we demonstrated in our previous study(Winstone et al., 2021), confirms that the FCS in Spike modulates IFITM entry restriction. Changes in the FCS would be predicted to increase the efficiency of viral fusion and entry at or near the plasma membrane, avoiding endosomal compartments where IFITMs 2 and 3 predominantly reside. Consistent with this, we show that the alpha Spike, as a PLV, is less sensitive to the cathepsin inhibitor E64D. Thus we propose that these changes in the alpha Spike have, in part, arisen to resist innate immunity.

At least two preprints suggest that variants of SARS-CoV-2 have begun to evolve further resistance to interferon-induced innate immunity (Guo et al., 2021; Thorne et al., 2021). In one, viral isolates over the pandemic show a reduced sensitivity to type I interferons in culture(Guo et al., 2021); in a second the alpha variant has a significantly reduced propensity to trigger pattern recognition in epithelial cells(Thorne et al., 2021). In contrast, another study shows no difference in IFN sensitivity of the new variants in African green monkey Vero-E6 cells(Michael Rajah et al., 2021), although species-specificity in viral sensitivity to ISGs is a well characterized trait. The SARS-CoV-2 genome contains multiple mechanisms to counteract host innate immune responses, and much remains to be learned about the mechanisms deployed by this virus and its relatives. While many reports on SARS-CoV-2 evolution have naturally focussed on the pressing concern of potential for vaccine escape, it is very unlikely that all selective adaptations that we see arising in VOCs can be solely due to escape from adaptive immunity. The alpha variant Spike, for example, only displays a minor reduction in sensitivity to neutralizing antibodies (NAbs) (Graham et al., 2021; Mahase, 2021; Planas et al., 2021; Shen et al., 2021). However, this VOC had a considerable transmission advantage, with suspicions that it may have arisen in an immunocompromised individual with a persistent infection giving ample time for changes to be selected that further evade innate immunity, including those that target viral entry(Corey et al., 2021; Kemp et al., 2021).

In terms of IFITM resistance of VOC Spike proteins, so far we have only seen marked change in phenotype for the alpha variant. This is despite the fact that delta, which has superseded alpha in many places around the world in 2021, also shows an adaptation for enhanced S1/S2 cleavage with an P681R change(Liu et al., 2021; Peacock et al., 2021b). This would suggest that efficient cleavage of S1/S2 is necessary but not sufficient for IFITM resistance, and indeed our data implicate a context dependency of the NTD deletion at 69/70. While unique to the alpha VOC, the 69/70 deletion has been observed in persistent infection of immunosuppressed individuals and is thought to enhance viral fitness and Spike stability (Meng et al., 2021). While deletions in the NTD do affect NAb binding, it is primarily the 144 deletion in the alpha Spike that escapes neutralization by NTD-directed Nabs (Chi et al., 2020) and we show that this has no impact on IFITM sensitivity. By contrast, the more pronounced antibody evasion by the beta, gamma and delta variants is related to mutations in the major neutralizing epitopes of the RBD, suggesting that they may well have been driven by antibody escape(Planas et al., 2021; Zhou et al., 2021). Viral glycoproteins are dynamic structures that shift through large-scale conformational changes while interacting with their cognate receptors mediating viral membrane fusion. Such context dependency is therefore likely to be complex and will arise under competing selective pressures. Indeed, we have previously shown that the HIV-1 envelope glycoprotein of transmitted viruses is IFITM insensitive and this contributes to their overall type I IFN resistance (Foster et al., 2016). As HIV-1 infection progresses over the first 6 months in an infected person, the circulating variants increase in IFN/IFITM sensitivity and this is determined by adaptive changes in Env that resist the early neutralizing antibody response (Fenton-May et al., 2013). Such escape has structural and functional implications for such dynamic proteins that may impact upon receptor interactions and route of entry into the target cell.

While furin cleavage of the SARS-CoV-2 Spike reduces its IFITM sensitivity, other interferon-induced proteins may contribute to this phenotype. The guanylate binding protein family, and particularly GBP2 and GBP5, have been shown to have a general antiviral activity against enveloped viruses by dysregulating furin processing of diverse viral and cellular proteins (Braun et al., 2019). Similarly, IFITM overexpression in HIV-infected cells can lead to their incorporation into virions and in some cases promote defects in glycoprotein incorporation (Tartour et al., 2014). Future studies will confirm whether either of these mechanisms are involved in the IFN-resistance associated with the P681H mutation in alpha.

In summary, the spike protein of SARS-CoV-2 alpha increases resistance to IFN-I and this correlates with the P681H mutation. Furthermore, this correlates with resistance to IFITM-mediated entry restriction. This suggests that in addition to adaptive immune escape, fixed mutations associated with VOCs may well also confer replication and/or transmission advantage through adaptation to resist innate immune mechanisms.

## MATERIALS AND METHODS

### Cells and plasmids

HEK293T-17 (ATCC, CRL-11268™), Calu-3 (ATCC, HTB-55™), A549-ACE2, Vero-E6, Vero-E6-TMPRSS2 and A549-ACE2 expressing the individual IFITM proteins were cultured in DMEM (Gibco) with 10% FBS (Invitrogen) and 200μg/ml Gentamicin (Sigma), and incubated at 37°C, 5% CO_2_. ACE2, TMPRSS2, and IFITM stable overexpression cells were generated as previously described (Winstone et al., 2021).

Codon optimised SARS-CoV-2 Wuhan Spike and ACE2 were kindly given by Dr. Nigel Temperton. Codon optimised variant Spikes (B.1.1.7, B.1.351) were kindly given by Dr. Katie Doores. Codon optimised variant Spikes (P1, B.617.2) were kindly given by Professor Wendy Barclay. Plasmid containing TMPRSS2 gene was kindly given by Dr. Caroline Goujon. Spike mutants were generated with Q5® Site-Directed Mutagenesis Kit (E0554) following the manufacturer’s instructions, and using the following forward and reverse primers:

D614G (GCTGTACCAGGGCGTGAATTGCA, ACGGCCACCTGATTGCTG) B.1.351. Δ242-244 (ATTTCATATCTTACACCAGGC, ATGCAGGGTCTGGAATCTG) D614G P681H (GACCAATAGCcacAGAAGAGCCAGAAGC, TGGGTCTGGTAGCTGGCG)

B117 ΔHRRA (AGAAGCGTGGCCAGCCAG, GCTATTGGTCTGGGTCTGGTAG) B117 H681P (GACCAATAGCcccAGAAGAGCCAG, TGGGTCTGGTAGCTGGCG) Δ144 (CATAAGAACAACAAGAGC, ATAAACACCCAGGAAAGG), N501Y (CCAGCCTACCtacGGCGTGGGCT, AAGCCGTAGCTCTGCAGAG), E484K (TAATGGCGTGAACGGCTTCAATTGCTACTT, CACGGTGTGCTGCCGGCC).

A549 stable cell lines expressing ACE2 (pMIGR1-puro), and IFITMs (pLHCX) were generated and selected as described previously (Winstone et al., 2021).

### Production of Pseudotyped Lentiviral Vectors (PLVs) and infection

HEK293T-17 were transfected with firefly luciferase expressing vector (CSXW), HIV gag-pol (8.91) and Spike plasmid with PEI-max as previously described (Winstone et al., 2021). Viral supernatant was then used to transduce each cell line of interest and readout measured by Luciferase activity 48 hours later (Promega Steady-Glo® (E2550)).

### Cyclosporin H assay

Cells were pre-treated with 30μM of Cyclosporin H (Sigma, SML1575) for 18 hours. Cells were then infected with PLVs and viral entry quantified by Luciferase activity 48 hours later.

### Passage and titration of SARS-CoV-2

PHE England strain 02/2020 was propagated in Vero-E6-TMPRSS2 cells and titre was determined by plaque assay (Winstone et al., 2021). Vero-E6-TMPRSS2 were infected with serial dilutions of SARS-CoV-2 for 1h. Subsequently, 2X overlay media (DMEM + 2% FBS + 0.1% agarose) was added, and infected cells were fixed 72 hours after infection and stained with Crystal Violet. Plaques were counted and multiplicity of infection calculated for subsequent experiments. A replication-competent alpha variant was kindly provided by Professor Wendy Barclay (Imperial College London)(Brown et al., 2021). All virus stocks were sequence confirmed in the Spike gene at each passage to ensure no loss of the FCS.

### Generation of recombinant full-length viruses

We used the previously described Transformation-Associated Recombination (TAR) in yeast method(Thi Nhu Thao et al., 2020), with some modifications, to generate the mutant viruses described in this study. Briefly, a set of overlapping cDNA fragments representing the entire genomes of SARS-CoV-2 Wuhan isolate (GenBank: MN908947.3) and the B.1.1.7 alpha variant were chemically synthesized and cloned into pUC57-Kan (Bio Basic Canada Inc and Genewiz, respectively). The cDNA fragment representing the 5’ terminus of the viral genome contained the bacteriophage T7 RNA polymerase promoter preceded by a short sequence stretch homologous to the *Xho*I-cut end of the TAR in yeast vector pEB2(Gaida et al., 2011). The fragment representing the 3’ terminus contained the T7 RNA polymerase termination sequences followed by a short segment homologous to the *Bam*HI-cut end of pEB2.

To generate Wuhan virus carrying the alpha variant spike, a mixture of the relevant synthetic cDNA fragments of the Wuhan and alpha variants was co-transformed with *Xho*I-*Bam*HI-cut pEB2 into the *Saccharomyces cerevisiae* strain TYC1 (MATa, ura3-52, leu2Δ1, cyh2^r^, containing a knockout of DNA Ligase 4) (Gaida et al., 2011) that had been made competent for DNA uptake using the LiCl_2_-based Yeast transformation kit (YEAST1-1KT, Merck). The transformed cells were plated on minimal synthetic defined (SD) agar medium lacking uracil (Ura) but containing 0.002% (w/v) cycloheximide to prevent selection of cells carrying the empty vector. Following incubation at 30^□^C for 4 to 5 days, colonies of the yeast transformants were screened by PCR using specific primers to identify those carrying plasmid with fully assembled genomes. Selected positive colonies were then expanded to grow in 200 ml SD-Ura dropout medium and the plasmid extracted. Approximately 4 μg of the extracted material was then used as template to *in vitro* synthesized viral genomic RNA transcripts using the Ribomax T7 RNA transcription Kit (Promega) and Ribo m7G Cap Analogue (Promega) as per the manufacturer’s protocol. Approximately 2.5 μg of the *in vitro* synthesized RNA was used to transfect ∼6 ×10^5^ BHK-hACE2-N cells stably expressing the SARS-CoV-2 N and the human ACE2 genes(Rihn et al., 2021) using the MessengerMax lipofection kit (Thermo Scientific) as per the manufacturer’s instructions. Cells were then incubated until signs of viral replication (syncytia formation) became visible (usually after 2-3 days), at which time the medium was collected (P0 stock) and used further as a source of rescued virus to infect VERO E6 cells to generate P1 and P2 stocks. Full genome sequences of viruses collected from from P0 and P1 stocks were obtained in order to confirm the presence of the desired mutations and exclude the presence of other spurious mutations. Viruses were sequenced using Oxford Nanopore as previously described(da Silva Filipe et al., 2021).

To generate Wuhan virus carrying alpha spike gene with the H681P mutation, we first introduced this mutation into the relevant alpha variant cDNA fragment by site-directed mutagenesis. This fragment was combined with those described above and the mixture was then used to generate plasmid pEB2 carrying the cDNA genome of Wuhan encoding the alpha spike H681P by the TAR in yeast procedure. The virus rescue and subsequent characterisation were performed as described above.

### Generation of Clinical Viral Isolates

Viruses were isolated on Vero.E6 cells (ATCC CRL 1586™) from combined naso-oropharyngeal swabs submitted for routine diagnostic testing by real-time RT-PCR and shown to be from the B.1.1.7 (alpha) variant by on-site whole-genome sequencing (Oxford Nanopore Technologies, Oxford, UK) (Pickering et al., 2021). Infected cells were cultured at 37°C and 5% CO2, in Dulbecco’s modified Eagle’s medium (DMEM, Gibco™, Thermo Fisher, UK) supplemented with 2% foetal bovine serum (FBS, Merck, Germany), pen/strep and amphotericin B.

All work performed with full-length SARS-CoV-2 preparations, as well as isolation and propagation of viral isolates from swabs, was conducted inside a class II microbiological safety cabinet in a biosafety level 3 (BSL3) facility at King’s College London.

### Infection with replication competent SARS-CoV-2

1.5×10^5^ A549-ACE2 cells were infected for 1 hour at 37°C with SARS-CoV-2 replication competent viruses at MOI 0.01 or 500 E gene mRNA copies/cell. 2×10^5^ Calu-3 cells were infected for 1h at 37°C with SARS-CoV-2 replication competent viruses at 5000 copies/cell. Media was replaced and cells were incubated for 48 hours at 37°C, after which cells or supernatant were harvested for RNA extraction or protein analysis.

### Interferon assays

Cells were treated with different doses of IFNβ (PBL Assay Science, 11415-1) for 18 hours prior infection. The following day media was replaced, and the infection performed as described above. Viral RNA levels in cells or supernatants were measured 48 hours after infection by RT-qPCR.

### siRNA knockdown of IFITM2

A549-ACE2 cells were reverse transfected using 20pmol of Non-targeting siRNA (D-001206-13-20) or IFITM2 siRNA (M-020103-02-0010) and 1μL of RNAi max (Invitrogen). Cells were incubated for 24h prior to a second round of reverse transfection. 8h later, cells were treated with different doses of IFNβ. Following 18h of IFN treatment cells were infected with full-length viruses as previously described.

### RT-qPCR

RNA from infected cells was extracted using QIAGEN RNeasy (QIAGEN RNeasy Mini Kit, 74106) following the manufacturer’s instructions. 1μL of each extracted RNA was used to performed one step RT-qPCR using TaqMan Fast Virus 1-Step Master Mix (Invitrogen). The relative quantities of envelope (E) gene were measured using SARS-CoV-2 (2019-nCoV) CDC qPCR Probe Assay (IDT DNA technologies). Relative quantities of E gene were normalised to GAPDH mRNA levels (Applied Bioscience, Hs99999905_m1).

Supernatant RNA was extracted using RNAdvance Viral XP (Beckman) following the manufacturer’s instructions. 5μL of each RNA was used for one-step RT-qPCR (TaqMan™ Fast Virus 1-Step Master Mix) to measured relative quantities of E and calibrated to a standard curve of E kindly provided by Professor Wendy Barclay.

### SDS-PAGE and Western blotting

Cellular samples were lysed in reducing Laemmli buffer at 95°C for 10 minutes. Supernatant or viral stock samples were centrifuged at 18,000 RCF through a 20% sucrose cushion for 1 hour at 4°C prior to lysis in reducing Laemmli buffer. Samples were separated on 8–16 % Mini-PROTEAN® TGX™ Precast gels (Bio-Rad) and transferred onto nitrocellulose membrane. Membranes were blocked in milk prior to detection with specific antibodies: 1:1000 ACE2 rabbit (Abcam, Ab108209),1:5000 GAPDH rabbit (Abcam, Ab9485), 1:5000 HSP90 mouse (Genetex, Gtx109753), 1:50 HIV-1 p24Gag mouse (48 ref before) 1:1000 Spike mouse (Genetex, Gtx632604), 1:1000 anti-SARS-CoV-2 N rabbit (GeneTex, GTX135357). Proteins were detected using LI-COR and ImageQuant LAS 4000 cameras.

### Ethics

Clinical samples were retrieved by the direct care team in the Directorate of Infection, at St Thomas Hospital, London, UK, and anonymised before sending to the King’s College London laboratories for virus isolation and propagation. Sample collection and studies were performed in accordance with the UK Policy Framework for Health and Social Care Research and with specific Research Ethics Committee approval (REC 20/SC/0310).

## ACKNOWLEDGEMENTS

We are grateful to Nigel Temperton, Caroline Goujon, Katie Doores, Wendy Barclay and Public Health England for reagents. We acknowledge the G2P-UK National Virology consortium funded by MRC/UKRI (grant ref: MR/W005611/1) and the Barclay Lab at Imperial College London for providing the alpha variant. We thank E. J. Louis, University of Leicester for generously providing the TAR in yeast system.

## AUTHORS CONTRIBUTION

Experiments were performed by MJL, HW, HDW and AD. SP, RPG, LS and GN collected, sequenced and isolated clinical viral isolates. MP, AHP, GDL, VMC, WF, NS, and RO generated reverse genetics-derived viruses. MJL, HW, HDW and AD analysed data. CMS provided reagents, funding support and advice. HW, MJL and SJDN analysed the data and wrote the manuscript. All authors edited the manuscript and provided comments.

## FIGURE LEGENDS

**Supplementary figure 1.**
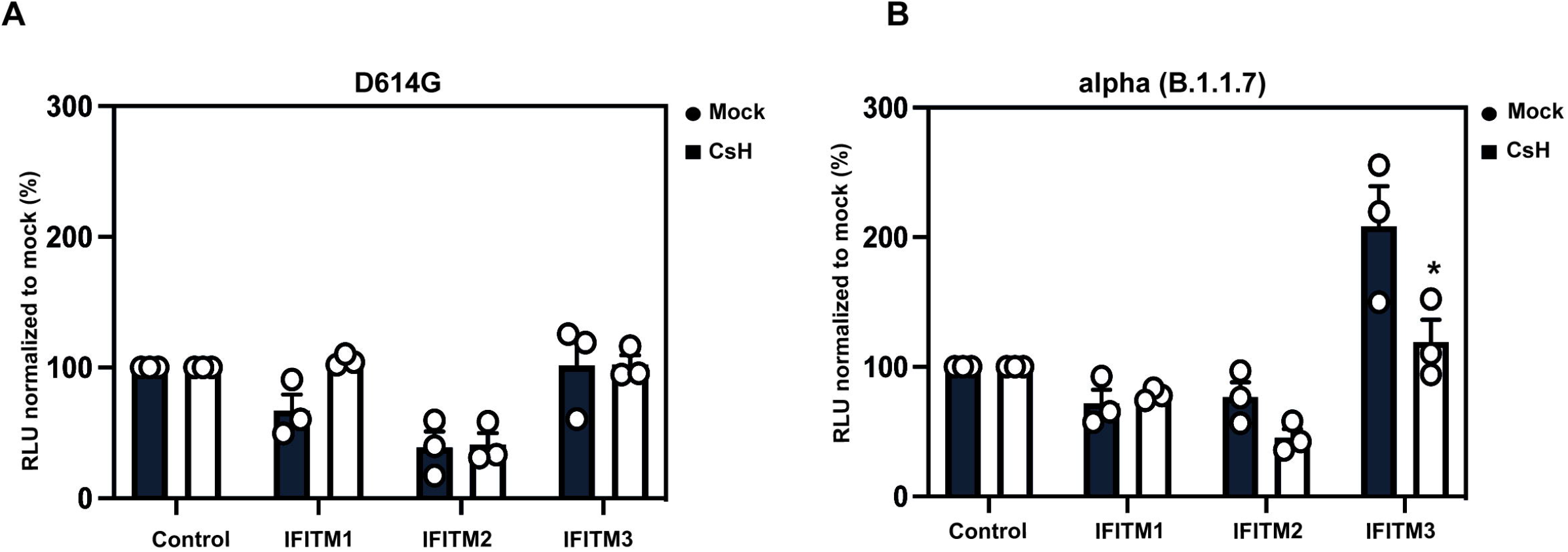
Cyclosporin H treatment abolishes IFITM3 enhancement of alpha PLVs. A) D614G PLVs pre-treated with Cyclosporin H. A549-ACE2s stably expressing the individual IFITMs were pre-treated with 30 μM of Cyclosporin H for 18 hours prior to infection with D614G PLVs. Infection was quantified by Luciferase activity 48 hours after infection and normalized to control cells. Data shown are mean ± SEM, n=3. Statistics were calculated in Prism using *t*-test (*P=<0.05). B) A549-ACE2s stably expressing the individual IFITMs were pre-treated with 30 μM of Cyclosporin H for 18 hours prior to infection with alpha PLVs. Infection was quantified by Luciferase activity 48 hours after infection and normalized to control cells. Data shown are mean ± SEM, n=3. Statistics were calculated in Prism using *t*-test, stars indiciate significance between IFITM3 mock and IFITM3 CsH (*P=<0.05).

**Supplementary figure 2.**
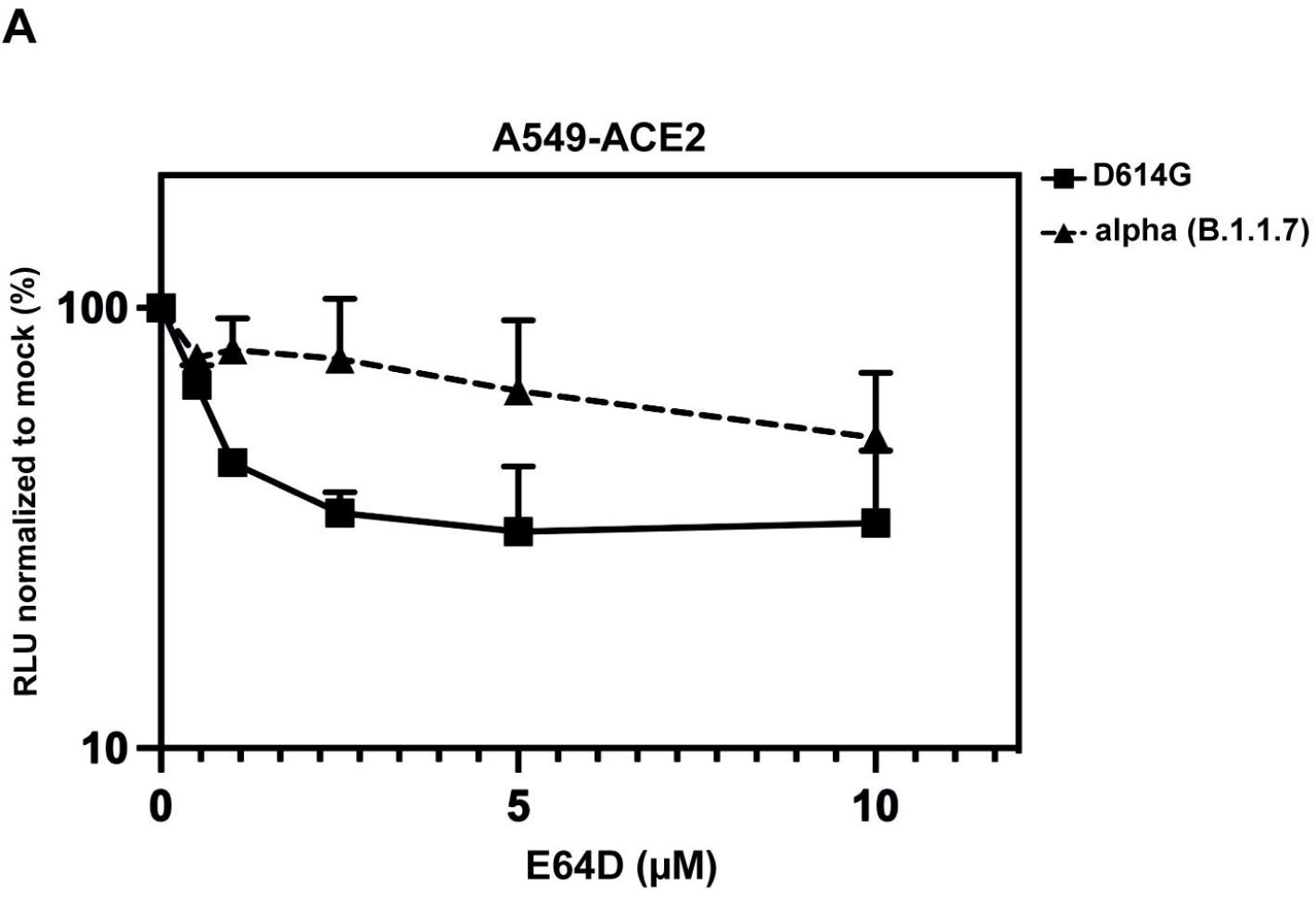
alpha PLVs are less sensitive to inhibition by E64D than D614G PLVs. A549-ACE2 cells were pre-treated with increasing concentrations of E64D for 1 hour prior infection with D614G or alpha PLVs. PLV entry was quantified by Luciferase activity 48 hours after infection and normalized to control cells. Data shown are mean ± SEM, n=3.

**Supplementary figure 3.**
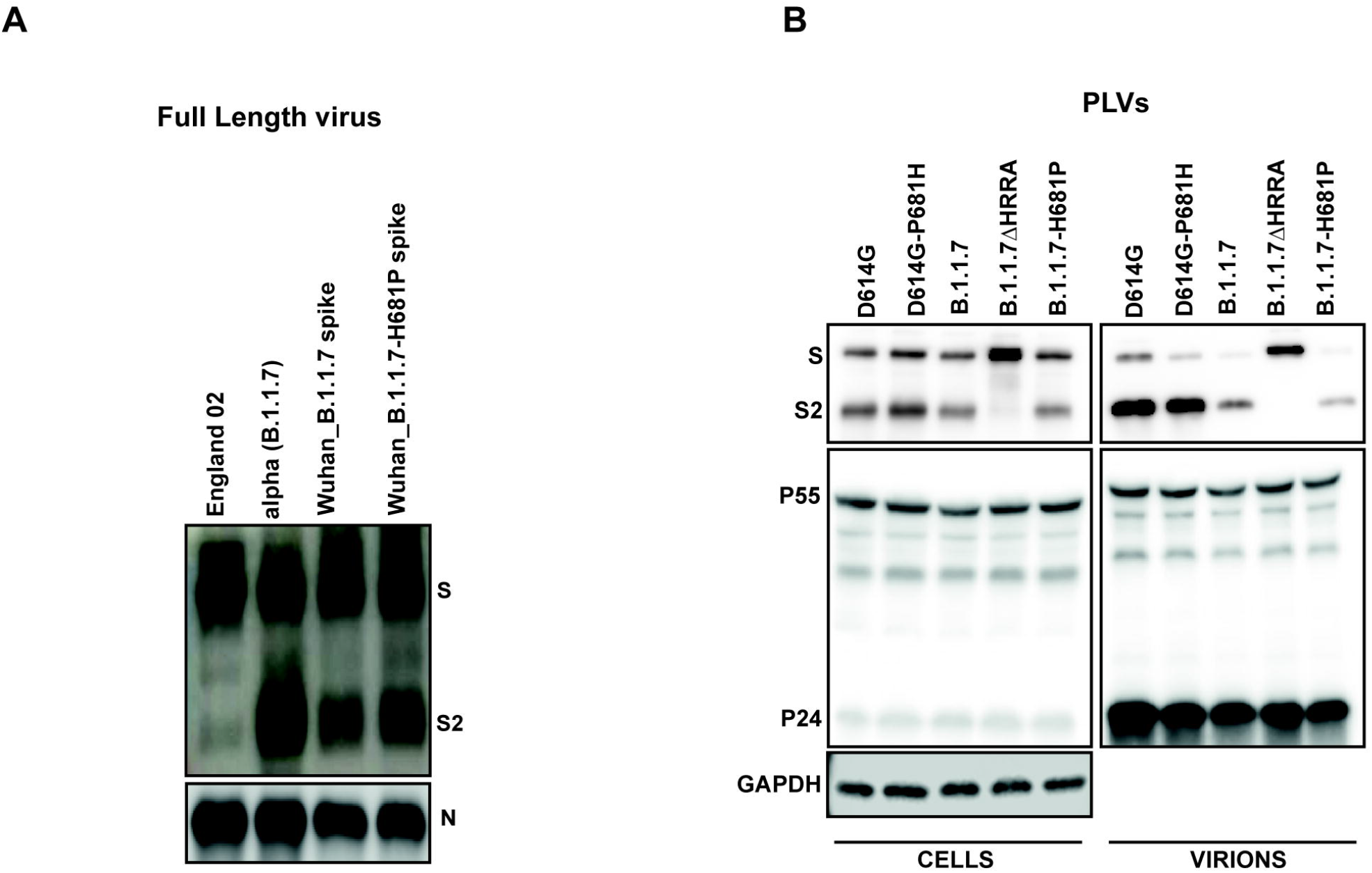
Spike processing of full-length virus and cleavage site PLVs mutants. A) England-02, alpha, Wuhan(B.1.1.7 Spike), Wuhan(B.1.1.7 Spike H681P) were purified through 20% sucrose and immunoblotted for Spike and N proteins. B) PLVs expressing different Spike mutants were produced in HEK293T-17 cells and cell lysates and supernatant immunoblotted for gag and Spike. Supernatant was purified through 20% sucrose.

**Supplementary figure 4.**
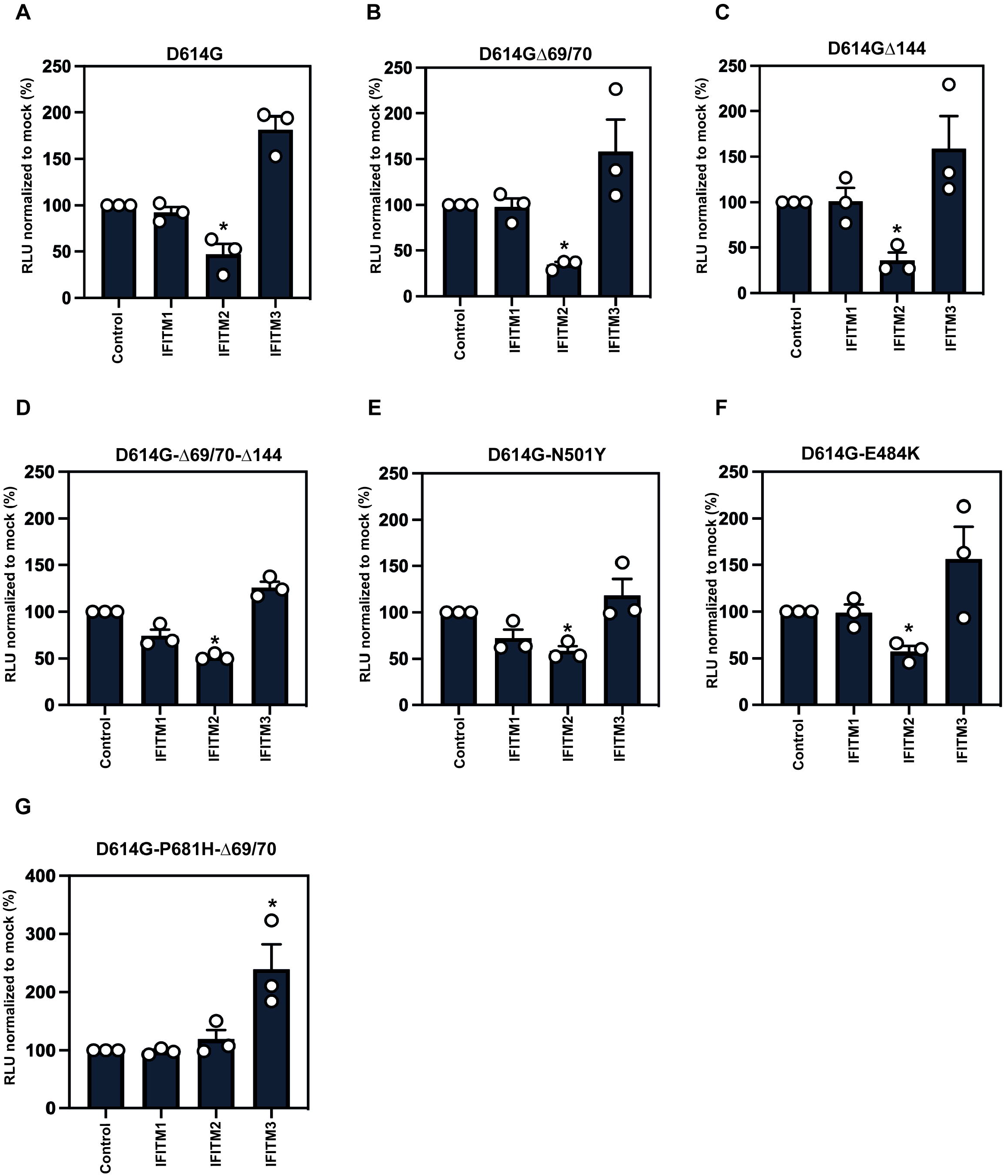
IFITM sensitivity of individual alpha Spike mutations. A–F) PLVs with individual alpha mutations were used to infect A549-ACE2 cells stably expressing the individual IFITMs. Infection was quantified by Luciferase activity 48 hours after infection and normalized to control cells. Data shown are mean ± SEM, n=3. Statistics were calculated in Prism using *t*-test, stars indiciate significance between each PLVs control cell and individual IFITM (*P=<0.05).

